# Lentiviral interleukin-10 gene therapy preserves fine motor circuitry and function after a cervical spinal cord injury in male and female mice

**DOI:** 10.1101/2020.06.05.137257

**Authors:** Jessica Y Chen, Emily J Fu, Paras R Patel, Hasan A Sawan, Kayla A Moss, Alexander J Hostetler, Sarah E Hocevar, Aileen J Anderson, Cynthia A Chestek, Lonnie D Shea

## Abstract

In mammals, spinal cord injuries often result in muscle paralysis through the apoptosis of lower motor neurons and denervation of neuromuscular junctions. Previous research shows that the inflammatory response to a spinal cord injury can cause additional tissue damage after the initial trauma. To modulate this inflammatory response, we delivered lentiviral anti-inflammatory interleukin-10, via loading onto an implantable biomaterial scaffold, into a left-sided hemisection at the C5 vertebra in mice. We hypothesized that improved behavioral outcomes associated with anti-inflammatory treatment are due to the sparing of fine motor circuit components. We examined behavioral recovery using a ladder beam, lower motor neuron apoptosis and muscle innervation using histology, and electromyogram recordings using intraspinal optogenetic stimulation at 2 weeks post-injury. Ladder beam analysis shows interleukin-10 treatment results in significant improvement of behavioral recovery at 2 and 12 weeks post-injury when compared to mice treated with a control virus. Histology shows interleukin-10 results in greater numbers of lower motor neurons and muscle innervation at 2 weeks post-injury. Furthermore, electromyogram recordings suggest that interleukin-10-treated animals have signal-to-noise ratios and peak-to-peak amplitudes more similar to that of uninjured controls than to that of control injured animals at 2 weeks post-injury. These data show that gene therapy using anti-inflammatory interleukin-10 can significantly reduce lower motor neuron loss, muscle denervation, and subsequent motor deficits after a spinal cord injury. Together, these results suggest that early modulation of the injury response can preserve muscle function with long-lasting benefits.

## 1 INTRODUCTION

The complexity of neural connections and inadequate nerve growth after a spinal cord injury (SCI) often results in permanent motor deficits or even complete paralysis [1]. Trauma can result in the apoptosis of lower motor neurons (LMNs), whose axons become detached from neuromuscular junctions (NMJs), leaving behind denervated motor endplates [2]. With sustained loss of neural input, these orphaned motor endplates will then release chemotactic signals to induce nearby, innervated NMJ axons to sprout growth cones [3, 4]. An orphaned motor endplate may then become re-innervated to form a functional NMJ. Alternatively, if the motor endplate remains denervated, it will disintegrate, the muscle fiber will atrophy, and paralysis will remain [5-9].

The early inflammatory response following trauma to the spinal cord can cause additional damage to neural tissues beyond the initial mechanical trauma [10, 11]. Immediately after the blood brain barrier is compromised, peripheral monocytes infiltrate and differentiate into macrophages that can exhibit phenotypes ranging from pro-inflammatory to anti-inflammatory. An inflammatory phenotype is highly linked to the secondary damage seen with an SCI, whereas the anti-inflammatory phenotype has been shown to encourage cell survival and tissue regeneration [12, 13]. Interleukin-10 (IL10) is an anti-inflammatory cytokine that is both expressed by macrophages and induces macrophage polarization towards an anti-inflammatory phenotype, which in turn results in a down-regulation of pro-apoptotic factors and an upregulation in anti-apoptotic factors [14-17]. In the absence of IL10, secondary damage is exacerbated [18]. Thus, shifting the post-injury response towards an anti-inflammatory phenotype has been a major target for therapeutic intervention.

The neuroprotective effects of IL10 treatment for an SCI have been investigated in several rodent studies. IL10 provides trophic support directly to neurons through the IL10 receptor [19], decreases post-injury inflammation [20], and increases myelination after a cervical SCI [21]. Many have shown that IL10 decreases lesion volume and apoptosis, while increasing behavioral recovery [20, 22-29]. However, the majority of animal SCI studies are carried out in female subjects [22-27, 29, 30], while most human SCIs occur in males [31]. Significant sex differences have been observed in immune responses after an SCI [32-34]. Furthermore, the majority of animal SCI studies are carried out at the thoracic level [23-29], while cervical level injuries are estimated to make up about half of all human SCIs [35]. In contrast to thoracic spinal tissues, the cervical enlargement contains a greater proportion of circuitry that is responsible for fine motor control, and the impact of IL10 treatment of cervical SCI has not been investigated.

In this study, we examined immunomodulation after a cervical SCI and assessed the sparing of fine motor circuitry and functional recovery, using both male and female mice. A left C5 lateral hemisection was performed, resulting in loss of function of the left arm. The resected tissue was replaced with a biomaterial bridge made of poly(lactide-co-glycolide) (PLG) loaded with lentivirus encoding for IL10, while lentiviral firefly luciferase (FLuc) served as a control. Lentivirus delivery has previously been shown to induce localized and sustained transgene expression [36]. Lentiviral IL10 expression can shift post-SCI gene expression towards an anti-inflammatory profile [37, 38]. We initially analyzed behavioral recovery using a ladder beam at 2 and 12 weeks post-injury (wpi). Then, using immunohistochemistry, we quantified LMNs, motor endplate densities, innervated NMJ densities, fraction of total NMJs that are innervated, and growth cone sprouting. We targeted the acromiotrapezius (ATZ) due to its topographical representation rostrally and throughout C5. In addition, we analyzed the flexor digitorum profundus & carpi ulnaris (FLX) together, due to their small size and topographical representation beginning in caudal C5 [39]. Finally, optogenetic stimulation was used to examine muscle activatability, which was quantified as a signal-to-noise ratio (SNR) and peak-to-peak (P2P) amplitude on an electromyogram (EMG). Our results show that early therapeutic intervention prevents some of the lifelong motor deficits by sparing the fine motor circuitry from secondary damage in both male and female mice.

## 2 MATERIALS & METHODS

### 2.1 Lentiviral Gene Delivery

Lentivirus containing pLenti-CMV-Luciferase or pLenti-CMV-IL10 was produced as previously described [20]. Briefly, HEK-293FT cells (American Type Culture Collection, Manassas, VA) were transfected with lentiviral packing vectors and plasmids of interest in Opti-MEM (Life Technologies, Carlsbad, CA, #31985-070) and Lipofectamine 2000 (Life Technologies, #11668-019). After 48hrs, the supernatant was collected and the virus was precipitated using PEG-It (Systems Biosciences, Palo Alto, CA, #LV825A-1) and stored at -80°C until use. Prior to surgery, 4e7 IFU of virus was loaded onto each PLG scaffold, which were fabricated as previously described using the gas foaming technique to fuse PLG particles into a matrix [20, 38, 40]. Using the particulate leaching technique, 63-106 µm NaCl served as sacrificial templates for pores to allow cell infiltration into the scaffold and a sugar mixture pulled into nine 150-250 µm diameter A-P parallel strands served as sacrificial templates for conduits to guide nerve growth [36, 41].

### 2.2 Animals and Surgery

All animal procedures conducted were approved by the Institutional Animal Care and Use Committee at the University of Michigan. For histology and ladder beam, mice used were 2-3-month-old C57BL/6 (Jackson Laboratories, Bar Harbor, ME, #000664) at the time of injury. For optogenetics, mice used were 2-3-month-old Thy1-ChR2-YFP line 18 (Jackson Laboratories #007612) at the time of injury. Uninjured data is derived from age-, sex-, and genotype-matched controls at the time of sacrifice.

Mice were anesthetized under 2% isoflurane. The surgery site was then shaved and disinfected with iodine and 70% ethanol. Bupivacaine (0.8ml/kg) was delivered locally to the incision site, and a C5 laminectomy was performed. Using a microfeather microscalpel (VWR, Radnor, PA, #72045-15), a left lateral hemisection of 1mm length (A-P) was performed at C5. The spinal tissue was excised and replaced with a lentivirus-loaded scaffold. Then the injury site was covered using GELFOAM (Pfizer, New York, NY), the overlying muscle was sutured using 5-0 Chromic Gut (Henry Schein, Melville, NY, #101-8824), and the skin was stapled. Mice received buprenorphine (0.1mg/kg) twice daily for 3 days, lactated ringer’s solution fluid supplement (5ml/100g) daily for 5 days, and enrofloxacin (2.5mg/kg) daily for 2 weeks.

### 2.3 Behavioral Analysis

C57BL/6 females only, due to persistent bladder issues in males, were trained to walk across a ladder beam consisting of 50+ rungs prior to surgery as previously described [42]. After surgery, at 2 and 12 wpi, animals were placed on the ladder beam and coerced to walk across while being recorded on video. A minimum of 3 trials was completed for each animal at each time point. The videos were randomized among a total of four individuals such that each video was quantified by two counters who were blinded to the treatment condition. Then each video’s quantifications were averaged between the two counters. A placement consists of 1) complete placement with all toes facing forward and the palm on the rung, 2) partial placements where the toes may be curled or the palm may not be centered though the limb is still weight-bearing, or 3) skipped rungs where the flanking rungs are placements. All others, such as misses or slips, were not counted. Examples can be seen in Fig. 1.

**Fig. 1.**
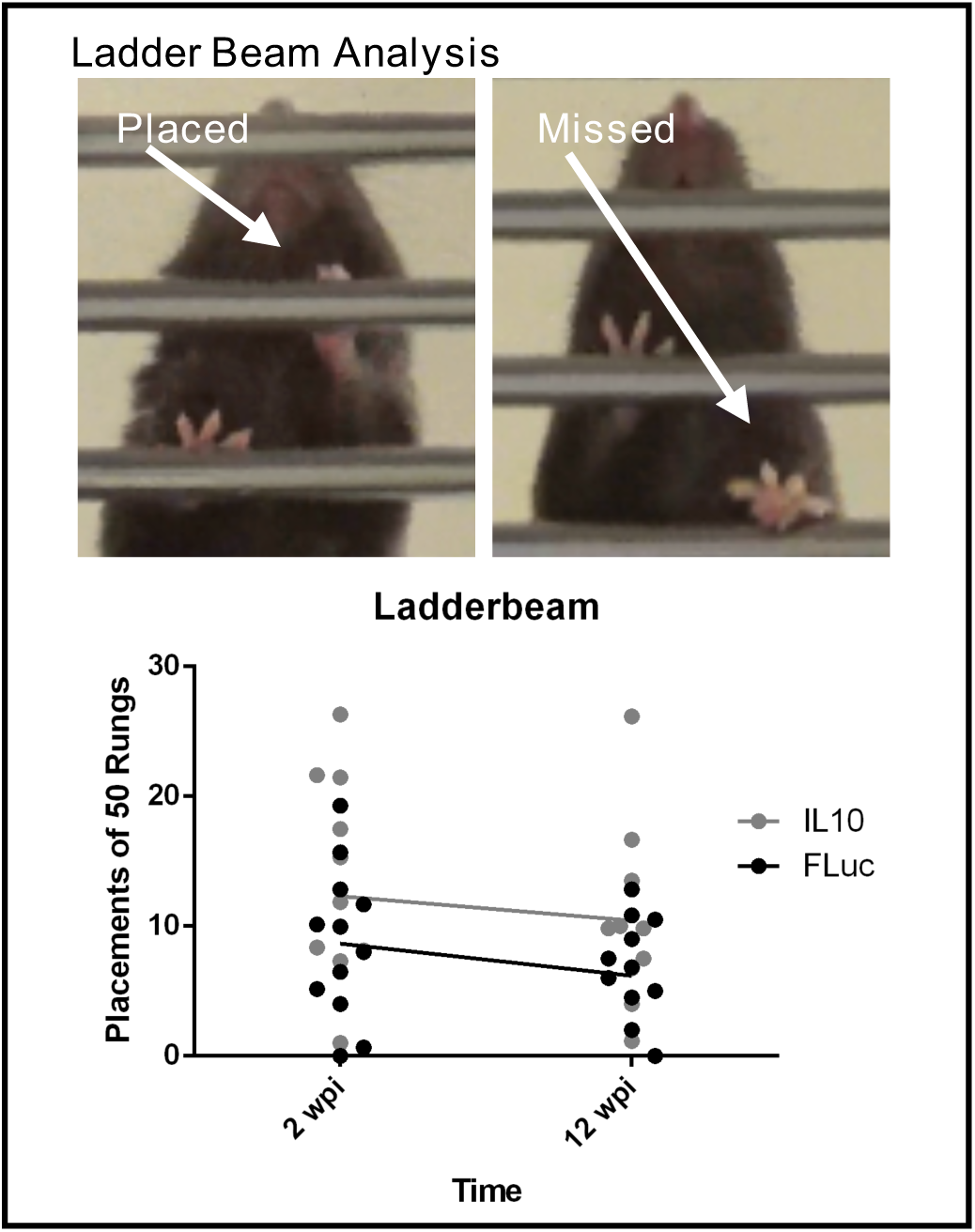
Paw placements (top) and counts (bottom) show IL10-treated animals perform significantly better when compared to FLuc-treated animals (Two-way ANOVA, n = 12-13, *p < 0.05 significant for virus condition)

### 2.4 Histological Analyses

At 2 wpi, spinal tissues and arm muscles were extracted, flash frozen in isopentane on dry ice, then cryosectioned at 12μm transverse and 14μm longitudinal, respectively. Spinal and muscle tissues were stained following a standard immunohistochemistry protocol with fixation and without fixation in 4% paraformaldehyde, respectively. Antibodies used were mouse anti-NeuN (1:250, Millipore, Burlington, MA, #MAB377), chicken anti-NF200 (1:250, AVES Labs, Davis, CA, #NFH), anti-bungarotoxin conjugated to Alexa Fluor 647 (1:500, Thermo Fisher, Waltham, MA, #B35450), and rabbit anti-GAP43 (1:100, Millipore, Burlington, MA, #AB5220). Secondary antibodies were Alexa Fluor 647 donkey anti-mouse (1:1000, Jackson ImmunoResearch, West Grove, PA #715-606-150), Alexa Fluor 488 goat anti-chicken (1:1000, Jackson ImmunoResearch, #103-547-008), and Alexa Fluor 594 goat anti-rabbit (1:1000, Jackson ImmunoResearch, #111-587-003).

Tissues were imaged using a Zeiss Axio Observer.Z1 microscope at 10X and processed in FIJI [43]. The analyze particles function (0.5-1 circularity, exclude on edges) in FIJI was utilized to count cells of 100-250 µm^2^ and 250-1100 µm^2^ on images that had a threshold established by a single observer who was blinded to the experimental condition. A minimum total of 300 neurons across a minimum of 4 spinal cord sections from each spinal cord sample was quantified. Total motor endplate density, innervated motor endplate density, fraction of innervated motor endplates, and fraction of motor endplates with a growth cone nearby were quantified and averaged across two counters who were blinded to the experimental condition. If an NF200+ or GAP43+ axon was colocalized or <1 axon diameter away from a motor endplate, then it would be considered innervated or in the process of being re-innervated, respectively. A minimum total of 150 motor endplates across a minimum of 3 muscle sections from each muscle sample was quantified.

### 2.5 Optogenetic Stimulation

The spinal column of Thy1-ChR2-YFP mice anesthetized under 2% isoflurane was exposed, and a C4-C6 laminectomy was performed from the dorsal side. The dura mater over C4-C6 was removed to allow insertion of a 1mm long LED probe (Plexon, Dallas, TX, #OPT/FS-Flat-110/125-1L) into the left C4, midway between the lateral edge of the spinal cord and the midline. Optical stimulation was delivered using a PlexBright Controller to drive a 465nm HELIOS headstage (Plexon, #OPT/LED_Blue_HELIOS_LC_Kit) in a faraday cage. The optical stimulation parameters were 20ms pulse width at 1Hz with a power of 0.04mW for a minimum of 3 minutes per animal.

### 2.6 EMG Data Acquisition and Processing

Digital out and reference signals from the PlexBright Controller were sent to a headstage (Tucker-Davis Technologies, Alachua, FL, #RA16AC) that was connected to a pre-amplifier (Tucker-Davis Technologies #RA16PA), which converted the inputted electrical signal into an outputted optical signal. To gain access to the muscle, small incisions were made in the skin and needle electrodes (Natus Neurology, Pleasanton, CA, #019-475400) were dually implanted into the left and right ATZ and FLX muscles. A single electrode in the left back ankle served as the reference signal. EMG and reference signals from the animal were connected to a second headstage and a second pre-amplifier. The optical outputs from each pre-amplifier were recorded by the same RX7 system (Tucker-Davis Technologies). A schematic for the equipment setup can be seen in Fig. 5a and an example EMG recording can be seen in Fig. 5b.

**Fig. 2.**
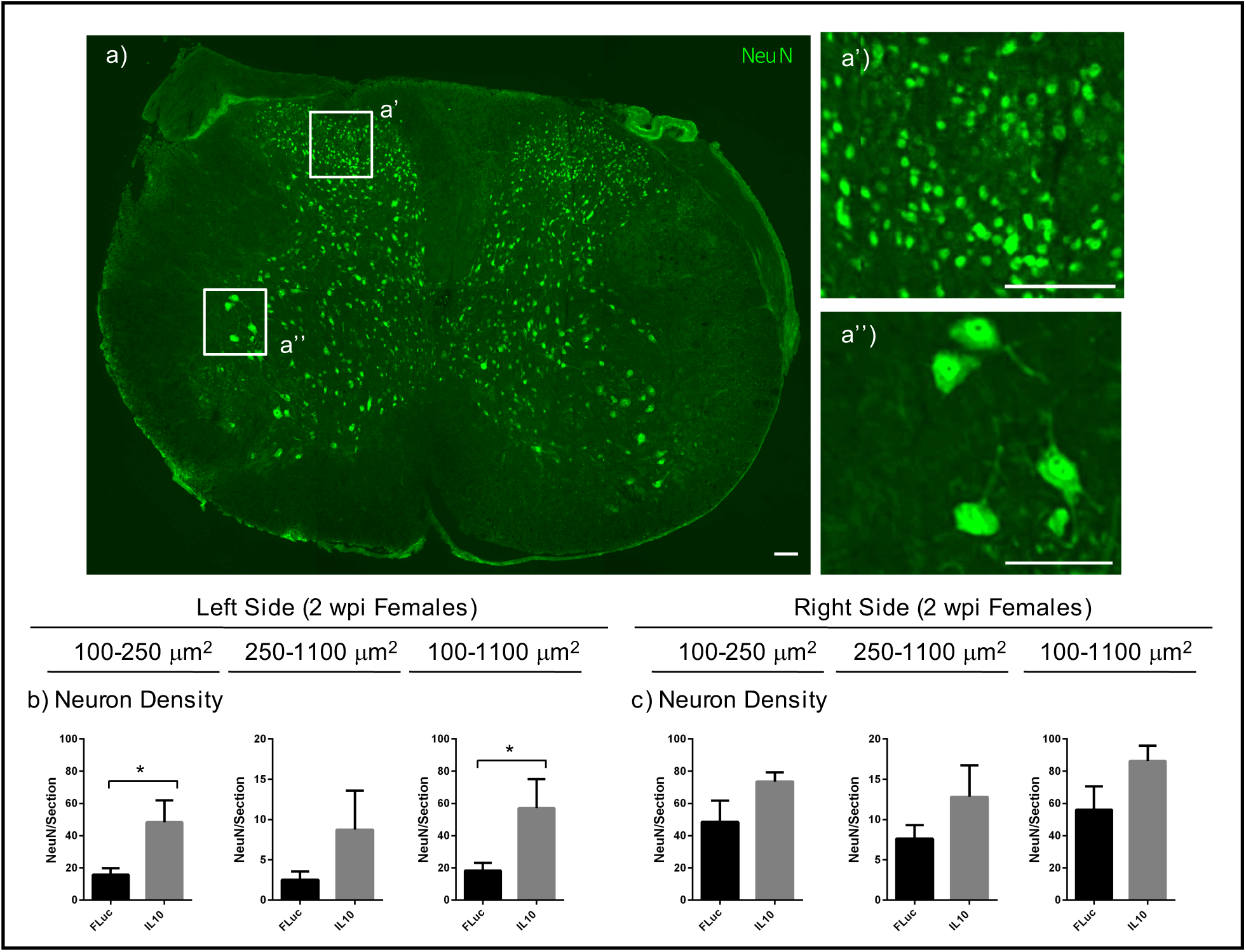
Tissue sections were stained for mature neurons (NeuN). Representative images of neurons per section (a), 100-250 μm^2^ size neurons (a’), and 250-1100 μm^2^ size neurons (a’’) are shown. Quantifications of neurons normalized to each section for the left (b) and right (c) side are shown for each size range for 2 wpi females. Error bars are ± SEM. n = 4 animals/condition. Mann-Whitney test *p < 0.05. Scale bar = 500 μm

**Fig. 3.**
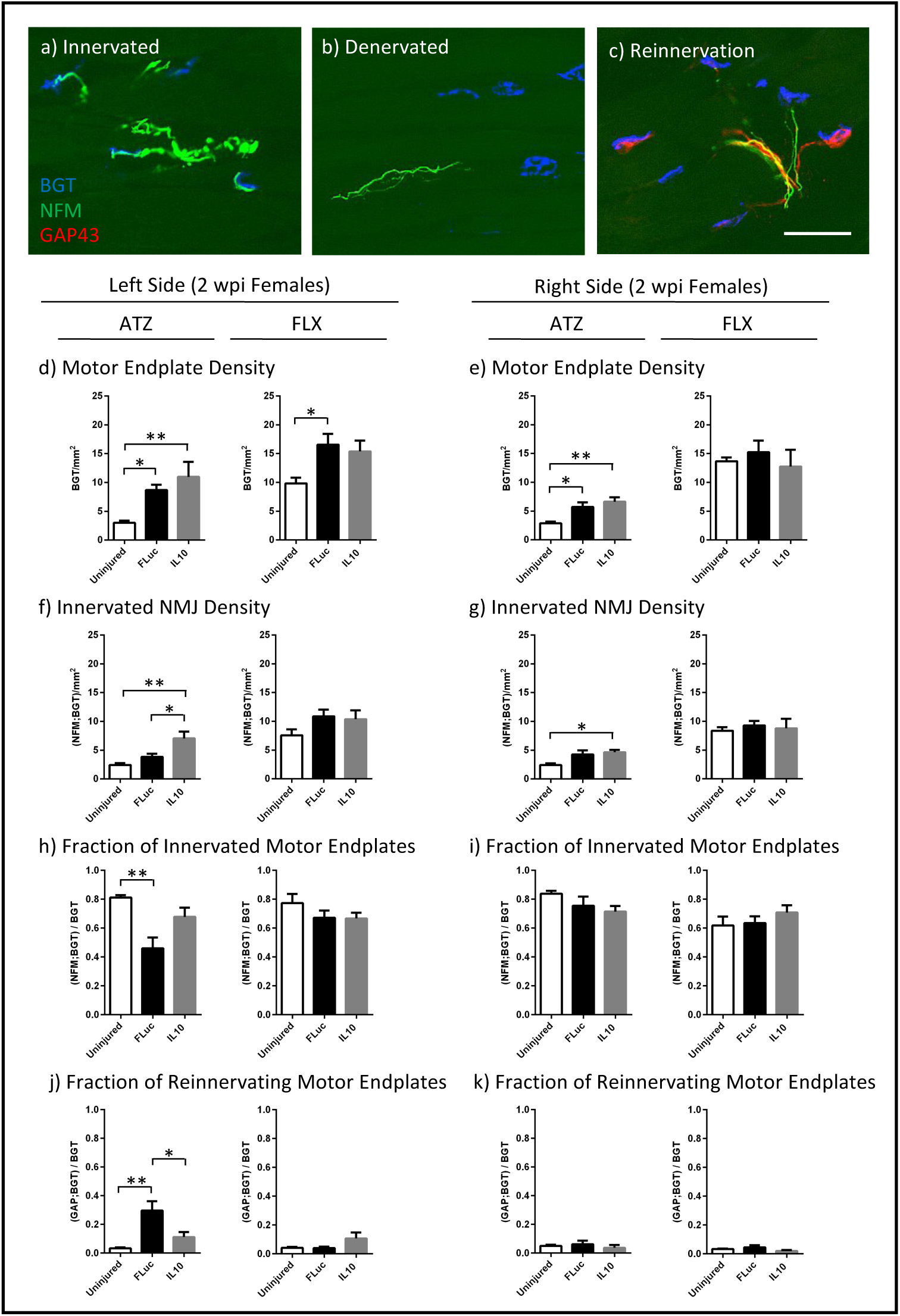
Tissue sections were stained for bungarotoxin (BGT) for motor endplates, neurofilament (NFM) for axons, and growth-associated protein 43 (GAP43) for growth cones. Representative images of NMJs that are (a) innervated, (b) denervated, or being (c) reinnervated are shown. Quantifications of motor endplate density normalized to muscle area (d, e), innervated NMJ density normalized to muscle area (f, g), fraction of total motor endplates that are innervated (h, i), and fraction of total motor endplates that are being reinnervated (j, k) are shown for 2 wpi females. Error bars are ± SEM. n = 5-6 animals/condition. One-Way ANOVA with Tukey Post-Hoc *p < 0.05, **p < 0.01. Scale bar = 50 μm

**Fig. 4.**
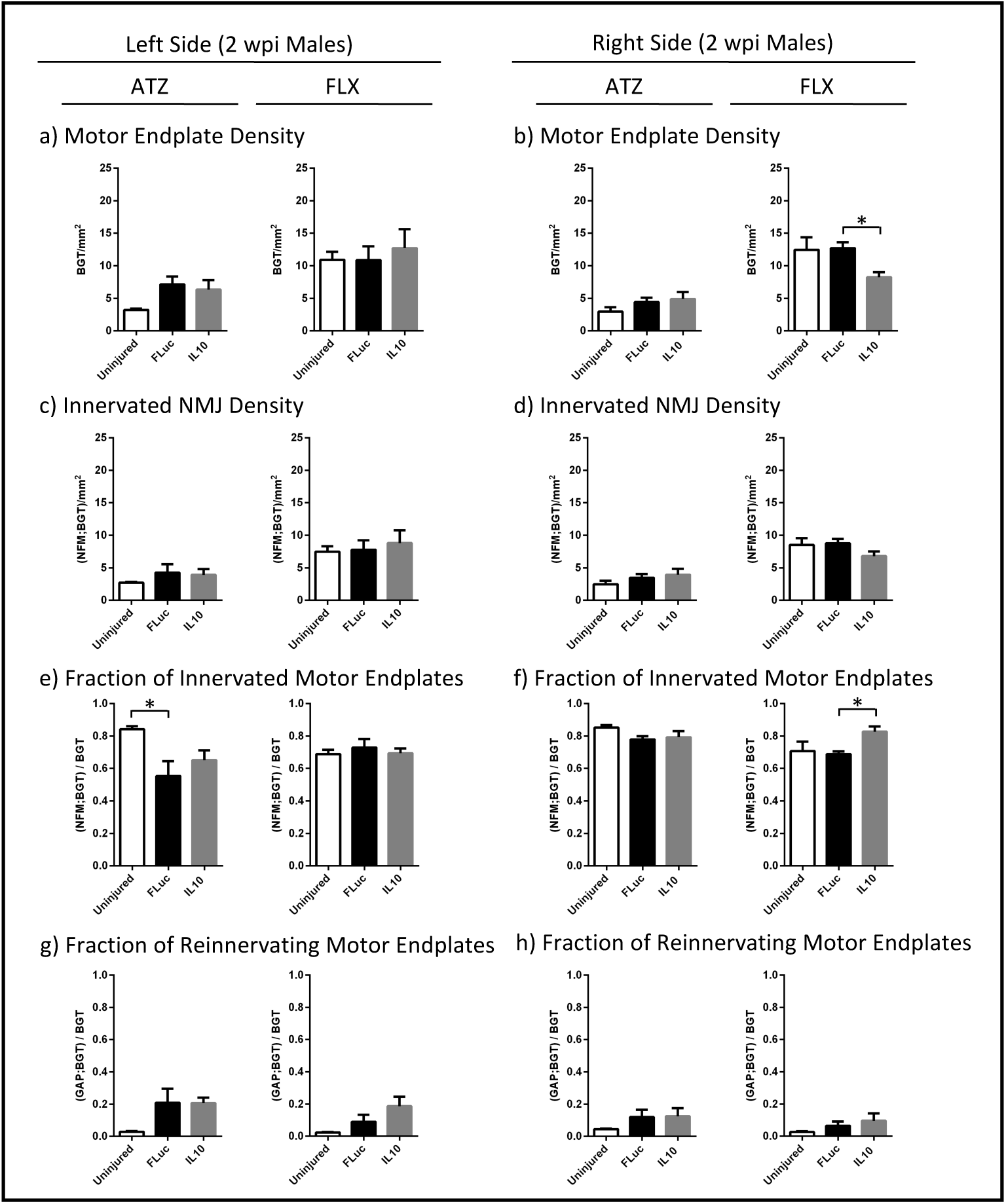
Quantifications of motor endplate density normalized to muscle area (a, b), innervated NMJ density normalized to muscle area (c, d), fraction of total motor endplates that are innervated (e, f), and fraction of total motor endplates that are being reinnervated (g, h) are shown for 2 wpi males. Error bars are ± SEM. n = 5-6 animals/condition. One-Way ANOVA with Tukey Post-Hoc *p < 0.05

**Fig. 5.**
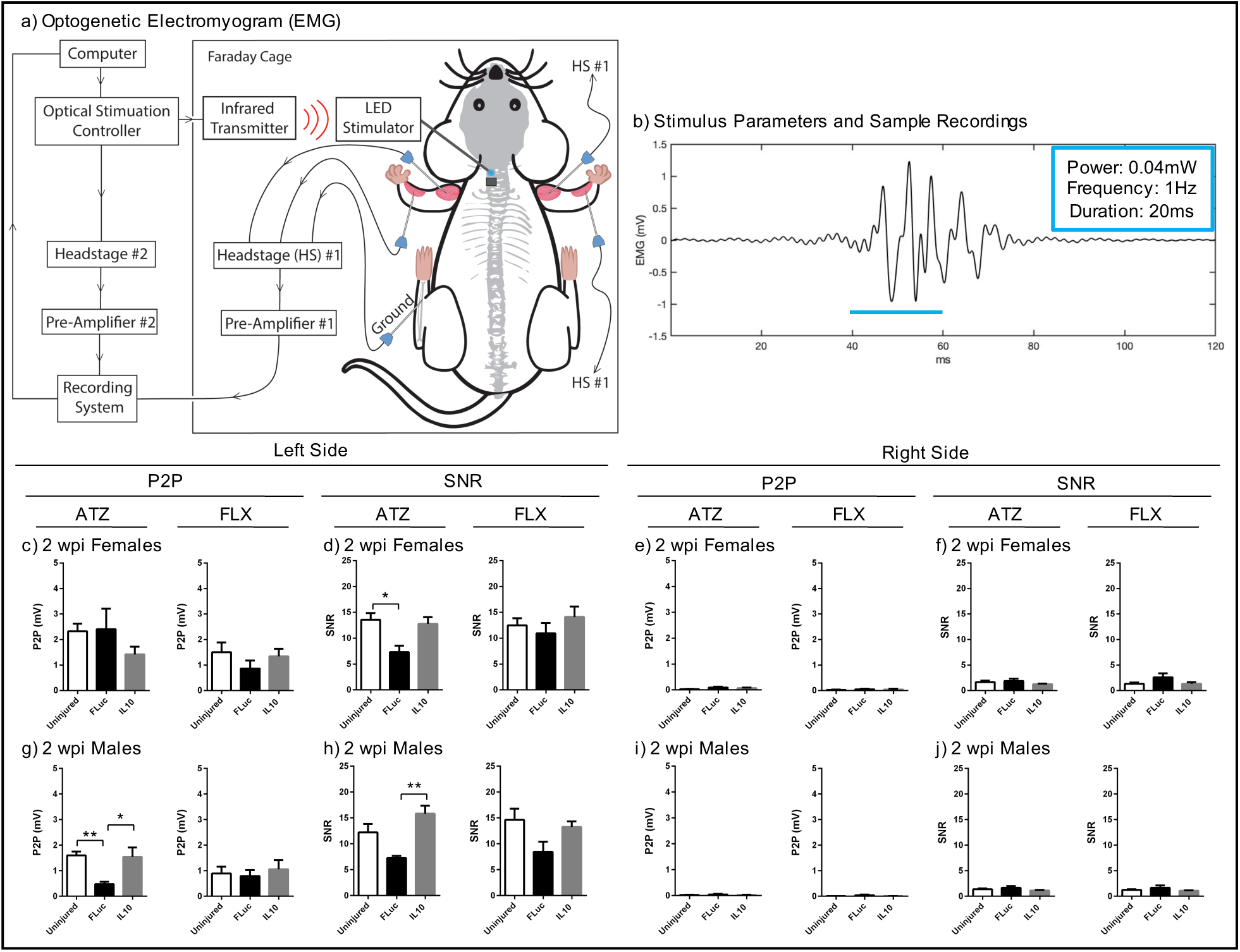
Schematic of the equipment setup (a) for optogenetically-evoked electromyogram recordings is shown, as well as the stimulus parameters (20ms pulses, 0.04mW, 3min, 1Hz) and a representative EMG trace (b). The peak-to-peak (c, e, g, i) and signal-to-noise ratios (d, f, h, j) for 2 wpi females (c, d, e, f), 2 wpi males (g, h, i, j) was quantified for the ATZ and FLX muscles. Error bars are ± SEM. n = 4-8 animals/condition. One-Way ANOVA with Tukey Post-Hoc *p < 0.05, **p < 0.01

Signals from both headstages were acquired at a synchronized rate of 24,414Hz with bandpass frequencies of 2.2Hz-7.5kHz. After acquisition, the data was converted to MATLAB® format using custom scripts. Next, the differential EMG electrode pairs were subtracted from each other and then band passed between 100-500Hz. To calculate the SNR for each muscle, a threshold was applied to the digital out signal representing the optical stimulation to determine the on and off times of the stimulation. Then, these times were used to calculate the root mean square values of the EMG signal during activation (stimulation on) and background (stimulation off). The ratio of the two values (on vs. off) was then used to calculate the SNR. To calculate the average P2P amplitude for each muscle, the maximum-to-minimum difference for each stimulation on time was determined, then averaged across all the stimulation on times for the length of that recording session.

## 3 RESULTS

### 3.1 Functional Recovery with IL10 Expression

Initial studies analyzed the behavioral response following a penetrating injury to the cervical spinal cord. A left-sided hemisection to the cervical enlargement at C5 allowed isolation of SCI-induced motor deficits to only the left forelimb. The injury level, injury sidedness, and difficulty of placing a weight-bearing limb onto a ladder beam rung allowed quantification of movement that requires significant fine motor control. Previous reports show uninjured control animals average 50 placements [42]. Animals receiving a bridge with a lentivirus encoding IL10 averaged 12.3 and 10.4 placements at 2 and 12 wpi, respectively. Animals receiving a bridge with a lentivirus encoding FLuc averaged 8.7 and 6.2 left forelimb placements at 2 and 12 wpi, respectively. A two-way ANOVA indicates that IL10-expression significantly increased placements relative to FLuc expression (p = 0.037), while no differences in performance were observed over time (p = 0.24) or for interaction between time and virus condition (p = 0.87) (Fig. 1).

### 3.2 Neuronal and NMJ Sparing with IL10

Inflammation following trauma can result in the apoptosis of neurons, and we thus quantified peri-injury neuronal density. NeuN+ neuron cell bodies range between 100-1100 µm^2^, with LMNs having larger sizes (250-1100 µm^2^) than interneurons (100-250 µm^2^) [44]. Histological analyses of spinal tissues at 2 wpi showed that animals treated with IL10 had significantly more interneurons and total neurons per section than animals treated with FLuc (Fig. 2). Since both the injured and contralateral side of FLuc-treated animals trended towards decreased neuron densities for all sizes of neurons analyzed, we added age- and sex-matched, uninjured controls for all subsequent experiments. All p-values are reported in Supplementary Table 1.

Next, we analyzed sparing in the peripheral nervous system through the quantification of motor endplate density and NMJ innervation status. The NMJs were characterized as innervated (overlap of NF200+ neurofilament (NFM) and bungarotoxin (BGT)) (Fig. 3a), denervated (separation of NFM and BGT) (Fig. 3b), and reinnervated (overlap of BGT and growth-associated protein 43 (GAP43)) (Fig. 3c). For both females (Fig. 3e, g, i, k) and males (Fig. 4b, d, f, h), a few significant differences were observed between conditions in the contralateral side, thereby supporting the use of an age- and sex-matched uninjured control. Additionally, for both female and male FLX muscles, the analysis of the NMJs largely showed no significant difference between the experimental conditions and uninjured controls. The lack of difference between injury and control suggests that the injury was consistently rostral to where FLX muscle innervation began.

In the left ATZ of females, a significant increase in motor endplate and innervated NMJ density per muscle area were observed in animals treated with either IL10 or FLuc relative to control uninjured animals (Fig. 3d, f). This result is consistent with previous reports showing that motor endplates will undergo fragmentation after a denervating event, resulting in an apparent increase in motor endplate density prior to the onset of muscle atrophy [45]. To account for fragmentation, we quantified the fraction of total motor endplates that are innervated and found that FLuc-treated animals had significantly decreased innervation when compared to that of control uninjured animals (Fig. 3h). Quantification of motor endplates colocalizing with growth cones showed that animals treated with FLuc also had the highest level of reinnervation (Fig. 3j). Since no difference in reinnervation or fraction of innervated motor endplates could be observed between IL10-treated and uninjured control animals, these data suggested IL10 spared more NMJs from denervation (Fig. 3h, j).

Overall, the differences observed for females (Fig. 3) were also seen in males, yet to a lesser extent (Fig. 4). The left ATZ in males treated with either IL10 or FLuc trended toward an increase in motor endplate and innervated NMJ densities (Fig. 4a, c). As with females, male mice treated with FLuc, but not IL10, had a significant decrease in their fraction of innervated motor endplates when compared to control uninjured animals (Fig. 4e). Animals treated with either IL10 or FLuc also trended towards an increase in their fraction of reinnervating motor endplates when compared to control uninjured animals (Fig. 4g). All p-values are reported in Supplementary Table 2.

### 3.3 Sparing of Muscle Activation with IL10

Next, we tested the hypothesis that NMJ differences identified by histology would influence muscle activation characteristics, which could be analyzed using electrophysiology. Denervation of an NMJ releases factors that can induce sprouting of nearby axons [3, 4]. If a denervated NMJ becomes captured by a new motor unit, then stimulation of the adopting motor unit will produce a greater contractile force that can be measured by an EMG as a compound action potential. If a denervated NMJ remains orphaned, stimulation of the motor unit that formerly owned that NMJ will produce a smaller compound action potential [5-9]. Therefore, we quantified the compound action potential as a P2P amplitude after optogenetic stimulation of channelrhodopsin positive C4 axons. Additionally, we quantified the SNR, because noise due to spontaneous fibrillations are a common characteristic of denervated muscles and can indicate damage to the motor circuitry independent of P2P changes.

No statistically significant differences in P2P and SNR values were observed between IL10-treated and uninjured control animals for both males and females, and for both the left ATZ and FLX muscles at 2 wpi (Fig. 5c, d, g, h), suggesting IL10 spared the motor circuitry from secondary damage. Meanwhile, females treated with FLuc had significantly lower left ATZ SNRs when compared to control uninjured animals (Fig. 5d), whereas males treated with FLuc had significantly lower left ATZ P2Ps and SNRs when compared to animals treated with IL10 (Fig. 5g, h). FLuc-treated males also had significantly lower left ATZ P2Ps when compared to uninjured control males (Fig. 5g). Together, these data show that an SCI without anti-inflammatory IL10 gene therapy, as is the case with animals treated with FLuc, has profound detrimental effects on muscle activation that can be identified in EMG recordings after optogenetic stimulation. Finally, no trends or statistically significant differences could be observed in the contralateral side (Fig. 5e, f, l, j), which all had low values, suggesting that the optogenetic stimulation light was contained within the left C4. All p-values are reported in Supplementary Table 3.

## 4 DISCUSSION

Our studies examined the potential for anti-inflammatory stimuli to enhance the fine motor circuitry and muscle activation in a cervical SCI model. The studies reported here validate that lentiviral IL10 gene therapy via a PLG implant into a left-sided, cervical SCI is associated with significantly improved performance of the left arm, as observed on a ladder beam. A number of groups have reported that IL10 treatment after a thoracic SCI can lead to intraspinal sparing of neurons, axons, and lesion size that is associated with improved locomotor function [23-29]. We have also previously demonstrated in a left-sided, thoracic SCI that IL10 can shift the inflammatory milieu toward a pro-regenerative phenotype, which was associated with significantly improved locomotor function [37, 38] and consistent with the results here in the cervical model. Thoracic injuries that employ behavioral tests such as the basso mouse scale for locomotion, which is controlled by central pattern generators (CPGs) that can be unilaterally activated to initiate bilateral locomotor behavior, may not fully capture the deficit from the unilateral injury. In contrast, cervical injuries are assessed through analysis of fine motor control and is controlled unilaterally [46]. Maintenance of fine motor control is also particularly relevant to humans, as we rely on forelimbs to a greater extent than rodents. Therefore, our studies provide further insight into IL10’s therapeutic benefits on the fine motor circuitry, in contrast to previous behavioral tests on hindlimb function in which CPGs and reflexes play a larger role.

Additionally, we demonstrate that IL10 expression within the spinal cord impacts processes in the periphery. Specifically, examination of muscles innervated at the level of injury indicated that IL10 decreased NMJ denervation and prevented the pathological increase in motor unit and spontaneous fibrillations in those same muscles. In mice, the ATZ is innervated by LMNs whose cell bodies reside between C2-C6 [39]. Within minutes after an injury, direct loss of the LMN cell body, such as removal via hemisection, can result in Wallerian degeneration of the axon and NMJ denervation [47, 48]. This first denervating event is likely unavoidable because the majority of hemisected LMN cell bodies from C5 were innervating the left ATZ. Later, peri-injury LMNs can undergo apoptosis that peaks at 3 days post injury in the highly inflammatory post-SCI environment, followed by axon degeneration and NMJ denervation [11, 49, 50]. This second denervating event is the target of our anti-inflammatory treatment. After a denervating event, motor endplates will undergo fragmentation [45, 51, 52]. The cumulative effects of direct removal of the LMN cell body and secondary damage due to inflammation can be seen most profoundly in the 2 wpi left ATZ of animals treated with FLuc, which had a significant increase in motor endplate density that reflects fragmentation and a significant decrease in the fraction of total motor endplates that are innervated when compared to uninjured control animals. Meanwhile, although the 2 wpi left ATZ of IL10-treated animals also showed significant motor endplate fragmentation, the fraction of total motor endplates that are innervated remained comparable to that of uninjured controls. Since animals treated with IL10 also had far fewer growth cones than animals treated with FLuc, suggesting a decreased need for reinnervation, together these data indicate a decrease in injury severity that is likely due to our anti-inflammatory treatment. We therefore identify IL10-induced sparing of NMJs in the left ATZ as a major contributor to improved fine motor control after a cervical SCI.

We report that IL10 treatment was effective in both male and female mice, though with some differences. All previous studies for delivery of IL10 used female rodents [21-27, 29, 37], except for one that did not indicate the sex of their animals [28]. Since differences in the neuroinflammatory response and locomotor recovery between male and female mice have been identified [53], IL10 treatment could potentially have disparate effects in each sex. For all the electrophysiological measures, no difference could be found between uninjured and IL10-treated animals, further supporting therapeutic benefits of IL10 in both sexes. Interestingly, control injury significantly reduced the P2P of the left ATZ, reflecting a decrease in motor unit size due to injury, in males but not in females. Since we found females treated with FLuc had significantly increased reinnervation, then reestablishment of a normal motor unit size might account for the apparently normal P2P seen in the left ATZ. Whether or not females have more or accelerated reinnervation relative to males remains to be determined. We do not anticipate that the normal P2P of the ATZ in females treated with FLuc reflects a lack of pathological outcomes, because increased pathological fibrillations explain why the SNR in females was decreased without a reduction in the P2P values. Additionally, while the electrophysiological effects of IL10 in males were clear, males showed fewer histological differences between conditions. Since all experiments were carried out in C57/BL6 background mice, and increased testosterone has been linked with decreased functional outcomes [54], we suspect that the observed sex differences reflect mechanistic differences in the injury response.

While tissue sparing is associated with preservation of function as measured by behavior and by optical stimulation at 2 wpi, no additional behavioral improvements were observed at 12 wpi. Hemisected intraspinal reflex and locomotor circuits will inevitably result in the loss of some synchronized muscle activation that is necessary for normal gait to occur. Additionally, increases in motor unit size due to reinnervation will decrease an animal’s control over their contractile force, such that the ratio of cognitive effort necessary to elicit a specific force will be shifted during fine motor movements [55]. If a muscle or neuron becomes inappropriately reinnervated after injury, the resulting synkinesis can further disrupt the synchronized muscle activation that is necessary for coordinated movement. Therefore, though sparing the motor circuitry by decreasing secondary damage due to inflammation can prevent some of the motor deficits that often results from an SCI, future studies might need to focus on the specificity of circuit reconnection in order for proper behavioral recovery to occur.

## 5 CONCLUSIONS

This study utilizes a left-sided, cervical level SCI to demonstrate that lentiviral IL10 can prevent some motor deficits from forming in the left arm. Intraspinal expression of IL10 can preserve NMJ innervation in the peripheral nervous system. Using optogenetic stimulation of C4 axons that originate from channelrhodopsin positive layer V neurons, we were able to examine muscle activation due solely to the fine motor circuitry and demonstrate that IL10 treatment can prevent pathological EMG signals in injury-affected muscles. Furthermore, these findings were mostly consistent in both male and female mice indicating potential applicability for both sexes. Collectively, our results indicate that early immunomodulatory intervention can yield long-term benefits.

## 6 ACKNOWLEDGEMENTS

The authors would like to thank Drs. Eva Feldman, Jonghyuck Park, Dominique Smith, and Courtney Dumont for their mentorship and guidance, as well as John Hayes and Rohit Maramraju for contributing their technical expertise. Funding for this work was provided by the National Institutes of Health (R01AI148076), the National Institute of Neurological Disorders and Stroke (U01NS094375 & UF1NS107659 for P.R.P.), the Office of the Director National institutes of Health (OT2OD024907 for P.R.P.), and the National Science Foundation (1707316 for P.R.P, and Graduate Research Fellowship for J.Y.C.), and the University of Michigan (Rackham Merit Followship for J.Y.C.).

---

**Supplementary Table 1.** Mann-Whitney test p-values associated with Fig. 2

| Figure # | Mann-Whitney p-value |
| --- | --- |
| 2b (100-250 $\mu\text{m}^2$) | 0.0286 |
| 2b (250-1100 $\mu\text{m}^2$) | 0.1714 |
| 2b (100-1100 $\mu\text{m}^2$) | 0.0286 |
| 2c (100-250 $\mu\text{m}^2$) | 0.1000 |
| 2c (250-100 $\mu\text{m}^2$) | 0.1000 |
| 2c (100-1100 $\mu\text{m}^2$) | 0.1000 |

**Supplementary Table 2.** One-Way ANOVA p-values and Tukey Post-Hoc p-values associated with Fig. 3 and 4

| Figure # | One-Way ANOVA p-value | Tukey Post-Hoc p-value |  |  |
| --- | --- | --- | --- | --- |
|  |  | Uninjured vs. Fluc | Uninjured vs. IL10 | Fluc vs. IL10 |
| 3d (ATZ) | 0.0099 | 0.0498 | 0.0093 | 0.5516 |
| 3d (FLX) | 0.0312 | 0.0318 | 0.0944 | 0.8703 |
| 3e (ATZ) | 0.0057 | 0.0262 | 0.0060 | 0.6114 |
| 3e (FLX) | 0.6898 | 0.8535 | 0.9502 | 0.6750 |
| 3f (ATZ) | 0.0034 | 0.4107 | 0.0031 | 0.0253 |
| 3f (FLX) | 0.1744 | 0.1788 | 0.3070 | 0.9542 |
| 3g (ATZ) | 0.0299 | 0.0715 | 0.0350 | 0.8618 |
| 3g (FLX) | 0.8300 | 0.8166 | 0.9627 | 0.9398 |
| 3h (ATZ) | 0.0040 | 0.0033 | 0.3289 | 0.0586 |
| 3h (FLX) | 0.3022 | 0.3645 | 0.3606 | 0.9972 |
| 3i (ATZ) | 0.2239 | 0.4378 | 0.2102 | 0.8188 |
| 3i (FLX) | 0.4727 | 0.9717 | 0.4871 | 0.5896 |
| 3j (ATZ) | 0.0037 | 0.0035 | 0.4920 | 0.0342 |
| 3j (FLX) | 0.1184 | 0.9991 | 0.1793 | 0.1465 |
| 3k (ATZ) | 0.6967 | 0.9157 | 0.8990 | 0.6726 |
| 3k (FLX) | 0.2711 | 0.6793 | 0.7029 | 0.2429 |
| 4a (ATZ) | 0.0981 | 0.0976 | 0.2081 | 0.8830 |
| 4a (FLX) | 0.8125 | > 0.9999 | 0.8545 | 0.8346 |
| 4b (ATZ) | 0.2882 | 0.4701 | 0.2771 | 0.9090 |
| 4b (FLX) | 0.0324 | 0.9885 | 0.0729 | 0.0445 |
| 4c (ATZ) | 0.5309 | 0.5249 | 0.6694 | 0.9653 |
| 4c (FLX) | 0.8221 | 0.9888 | 0.8263 | 0.8858 |
| 4d (ATZ) | 0.3750 | 0.5983 | 0.3528 | 0.8872 |
| 4d (FLX) | 0.2022 | 0.9758 | 0.3373 | 0.2200 |
| 4e (ATZ) | 0.0334 | 0.0277 | 0.1648 | 0.5640 |
| 4e (FLX) | 0.7407 | 0.7623 | 0.9936 | 0.8060 |
| 4f (ATZ) | 0.1895 | 0.1913 | 0.3198 | 0.9313 |
| 4f (FLX) | 0.0357 | 0.9287 | 0.1008 | 0.0411 |
| 4g (ATZ) | 0.0808 | 0.1106 | 0.1153 | 0.9997 |
| 4g (FLX) | 0.0689 | 0.5723 | 0.0599 | 0.2928 |
| 4h (ATZ) | 0.3778 | 0.4569 | 0.4137 | 0.9961 |
| 4h (FLX) | 0.3576 | 0.6860 | 0.3260 | 0.7781 |

**Supplementary Table 3.** One-Way ANOVA p-values and Tukey Post-Hoc p-values associated with Fig. 5

| Figure # | One-Way ANOVA p-value | Tukey Post-Hoc p-value |  |  |
| --- | --- | --- | --- | --- |
|  |  | Uninjured vs. Fluc | Uninjured vs. IL10 | Fluc vs. IL10 |
| 5c (ATZ) | 0.2207 | 0.9914 | 0.2591 | 0.3220 |
| 5c (FLX) | 0.5117 | 0.4834 | 0.9390 | 0.6881 |
| 5d (ATZ) | 0.0207 | 0.0193 | 0.9010 | 0.0532 |
| 5d (FLX) | 0.5296 | 0.8287 | 0.7701 | 0.5060 |
| 5e (ATZ) | 0.1146 | 0.1129 | 0.3678 | 0.6649 |
| 5e (FLX) | 0.7058 | 0.7199 | 0.8262 | 0.9674 |
| 5f (ATZ) | 0.4284 | 0.8900 | 0.5942 | 0.4348 |
| 5f (FLX) | 0.1101 | 0.1239 | > 0.9999 | 0.1492 |
| 5g (ATZ) | 0.0052 | 0.0066 | 0.9763 | 0.0162 |
| 5g (FLX) | 0.8323 | 0.9659 | 0.9160 | 0.8206 |
| 5h (ATZ) | 0.0045 | 0.0607 | 0.1991 | 0.0035 |
| 5h (FLX) | 0.1001 | 0.0917 | 0.8698 | 0.2602 |
| 5i (ATZ) | 0.4795 | 0.5936 | 0.9606 | 0.4900 |
| 5i (FLX) | 0.0545 | 0.0685 | > 0.9999 | 0.0961 |
| 5j (ATZ) | 0.2971 | 0.6497 | 0.6737 | 0.2670 |
| 5j (FLX) | 0.2472 | 0.4042 | 0.8741 | 0.2440 |

